# Interaction between histone residue H3K56 and mismatch repair protein MutSβ drives CAG repeat expansion

**DOI:** 10.1101/2024.09.29.615664

**Authors:** Jinzhen Guo, Wei Yang, Guo-Min Li

**Author notes:** Correspondence to: Wei Yang, Laboratory of Molecular Biology, National Institute of Diabetes and Digestive and Kidney Diseases, NIH, Bethesda, MD 20892; Guo-Min Li, Institute for Cancer Research, Chinese Institutes for Medical Research, Beijing, China 100069.

## Abstract

DNA repair protein MutSβ promotes CAG•CTG repeat expansions, which cause ∼20 untreatable neurodegenerative disorders including Huntington’s disease (HD). However, how MutSβ is recruited onto chromatin to enable repeat expansion is unknown. Here, we show that H3K56 interacts with MutSβ and brings it to chromatin. The H3K56-MutSβ interaction is regulated by H3K56 acetylation (H3K56ac) conditions, with acetylation inhibiting but deacetylation facilitating MutSβ chromatin recruitment. Blocking the H3K56-MutSβ interaction by either disrupting the MutSβ aromatic packet or increasing the H3K56ac level by removing histone deacetylases stabilizes CAG repeats, while depleting histone acetyltransferase p300/CBP promotes the H3K56-MutSβ interaction and CAG•CTG repeat instability, including expansion of CAG repeats of the huntingtin gene. Therefore, our study suggests a novel mechanism, by which MutSβ induces triplet repeat expansion, and new strategies of blocking the H3K56-MutSβ interaction to prevent HD and other triplet repeat expansion-caused disorders.

## Introduction

Trinucleotide repeat (TNR) expansions cause over 30 severe human neurodegenerative disorders, including Huntington’s disease (HD), myotonic dystrophy type 1 and fragile X syndrome ^1–5^. These diseases are currently untreatable, mainly because the mechanism of expansion is not fully understood. DNA mismatch repair (MMR) is well-known for its genome maintenance role by correcting mismatches generated during DNA replication ^6–9^. The MMR system relies on MutSα (a MSH2-MSH6 heterodimer) and MutSβ (a MSH2-MSH3 complex) to preferentially identify base-base mismatches and small insertion-deletion loops, respectively. While MutSα is a critical factor in genome maintenance, MutSβ has surprisingly been identified as a major driver of TNR expansions in HD patients and model systems ^10–16^. Thus, removing MutSβ or blocking its DNA-binding or ATPase activity is considered the effective treatment for HD and other TNR expansion-caused diseases ^17,18^. However, this approach is likely problematic, as MutSα exhibits almost identical structure and enzymatic activities of MutSβ ^19–22^.

It is known that MutSα is located onto chromatin by physically interacting with the histone mark H3K36me3 (trimethylated H3 lysine 36) through its MSH6 PWWP domain ^23^. Disrupting the H3K36me3-MutSα interaction results in loss of MMR function and a hypermutable phenotype ^23–25^. Since the MSH3 subunit of MutSβ does not have a PWWP domain, MutSβ chromatin recruitment has to rely on a non-H3K36me3 histone mark. Identifying this histone mark will provide opportunities to block the MutSβ function in repeat expansions. Despite extensive efforts, the histone mark that recruits MutSβ and the MutSβ domain responsible for interacting with chromatin remain unknown.

Interestingly, histone modifications, particularly acetylation and deacetylation, have been linked to HD. First, histone deacetylases (HDACs) stimulate CAG repeat expansion ^13,26^, and this stimulation is suppressed by HDAC inhibitors or gene knockouts ^27–31^. Second, depleting histone acetyltransferases (HATs), e.g., p300/CBP in mammals and Rtt109 in yeast, promotes TNR expansions ^26,32,33^. However, these activities have been thought to mediate TNR expansions by regulating the expression of targeted genes or acetylation/deacetylation of key proteins including MutSβ ^31,34–36^. We hypothesize that these HATs and HDACs modulate CAG repeat instability by controlling the acetylation level of a histone lysine residue that recruits MutSβ to chromatin.

Here, we show that this lysine residue is H3K56, which recruits MutSβ onto chromatin by specifically interacting with an MSH3 aromatic packet. The H3K56-MutSβ interaction is regulated by the H3K56 acetylation status, with acetylation inhibiting but deacetylation facilitating the interaction. Blocking the H3K56-MutSβ interaction by either disrupting the MutSβ aromatic packet or acetylating H3K56 stabilizes CAG repeats, while deacetylating H3K56 stimulates CAG repeat instability. Therefore, our results not only elucidate the molecular mechanism by which histone deacetylation stimulates CAG repeat expansions, but also provide new strategies to develop therapeutic drugs for HD and other CAG•CTG repeat expansion diseases by disrupting the H3K56-MutSβ interaction.

## Results

### MutSβ interacts with H3K56

To identify the specific histone H3 mark responsible for recruiting MutSβ, we performed protein pulldown experiments using purified MutSβ and various biotinylated H3 N-terminal peptides with or without a methylated or acetylated lysine residue (Figure 1**a**). This is based on the fact that methylation of H3K36 is responsible for recruiting DNA repair enzymes ^23,37,38^ and that histone acetylation status modulates CAG repeat stability ^13,26–31^. As shown in Figure 1**b**, the peptide containing H3 residues 44-63 with an unmodified H3K56 pulled down MutSβ (lane 4), but the one containing the same H3 residues with an acetylated H3K56 (H3K56ac, lane 7) or methylated H3K56 (H3K56me1, lane 8) failed to do so, nor do those containing a modified or unmodified H3K9 (lanes 2 and 5) or H3K36 (lanes 3 and 6). These results indicate that it is H3K56 that interacts with MutSβ and not H3K9 or H3K36; and that the H3-MutSβ interaction depends on an unmodified H3K56, while any modifications on H3K56 abolish the interaction. To confirm this result, we conducted coimmunoprecipitation (Co-IP) assays in HeLa nuclear extracts using an H3 antibody in the presence or absence of a competitive or non-competitive H3 peptide. The results revealed that MSH3 (i.e., MutSβ) could be coimmunoprecipitated by the H3 antibody in the presence of an H3K56-acetylated peptide (Figure 1**c**, lane 4) or no H3 peptide (Figure 1**c**, lanes 1 and 2), but the MSH3 coprecipitation was blocked when a K56-unmodified H3 peptide was added to the Co-IP reaction (Figure 1**c**, lane 3). The blockage of the endogenous MutSβ-H3K56 interaction by the H3K56-unmodified peptide strongly suggests a specific interaction between H3K56 and MutSβ. We also noted that addition of the peptide with H3K56ac in the Co-IP reaction slightly enhanced the MSH3 pulldown (Figure 1**c**, compare lane 4 with lanes 1 and 2), which is probably due to H3K56ac’s ability to compete with the natively unmodified H3K56 for other histone binding proteins, making the natively unmodified H3K56 available for MutSβ binding.

**Fig. 1.**
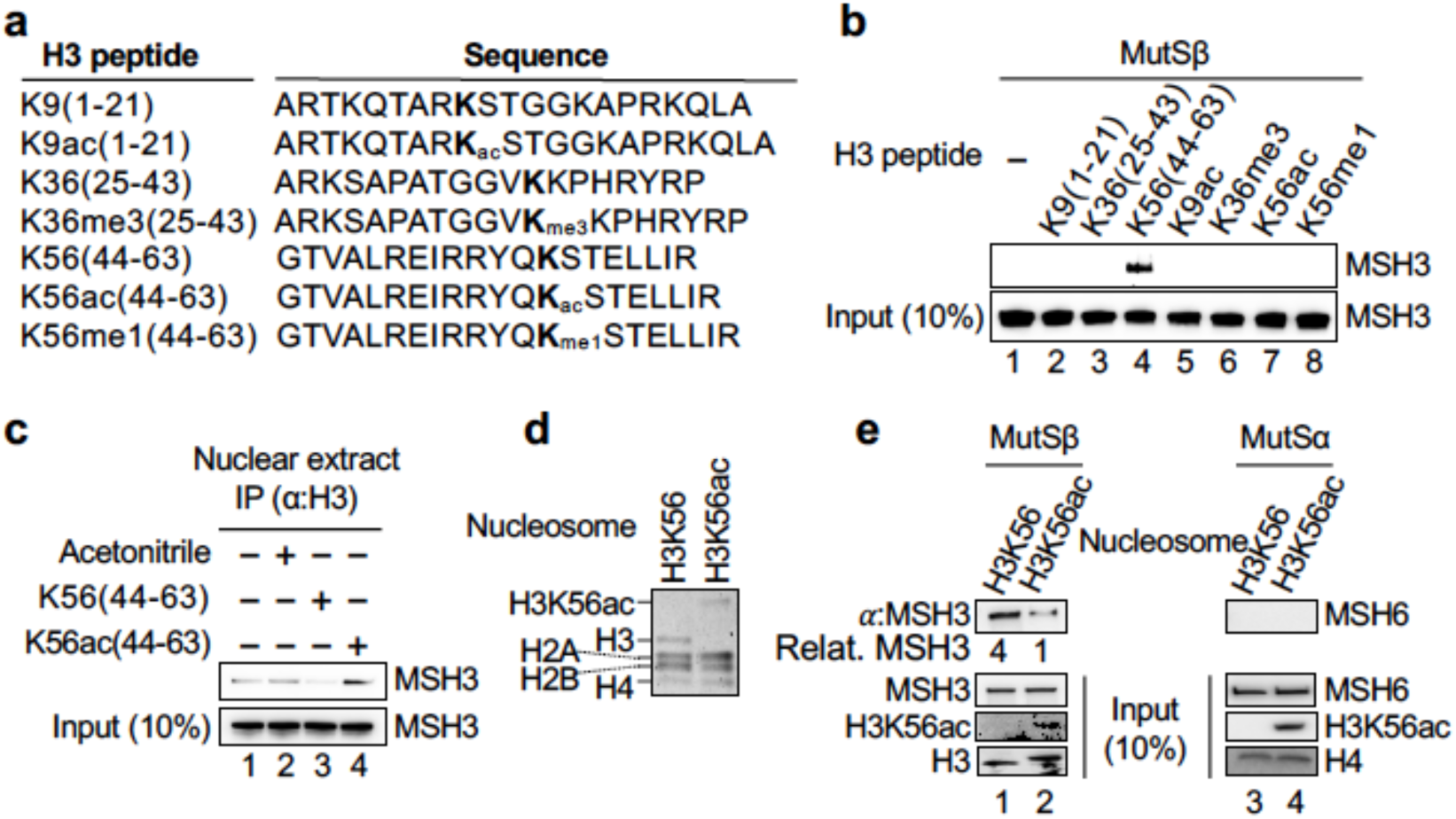
MutSβ interacts with H3K56 *in vitro*. **a,** Biotinylated H3 N-terminal peptides. **b,** Peptide pulldown assay to determine specific interaction between MutSβ and H3K56. Purified MutSβ was incubated with various biotinylated H3 peptides and streptavidin-conjugated beads, as indicated, and the protein pulled down was detected by an MSH3 antibody. **c,** Co-IP assays to determine specific interaction between MutSβ and H3K56 in HeLa nuclear extracts using an H3 antibody in the presence of the peptide resolvent acetonitrile or a peptide competitor K56(44-63) or K56ac(44-63), as indicated. The precipitates were fractionated in SDS-PAGE, followed by Western blotting using an MSH3 antibody. **d,** SDS-PAGE gel showing protein components of reconstituted nucleosomes. **e,** Nucleosome pulldown assay showing specific interaction between MutSβ (but not MutSα) with the H3K56-containing nucleosome but not the H3K56ac-containing nucleosome.

We next determined whether MutSβ interacts with H3K56 on nucleosome, the basic unit of chromatin. We reconstituted mononucleosomes with unmodified H3K56 or H3K56ac using the biotinylated nucleosome positioning sequence 601 ^39^ and purified recombinant histone proteins (Figure 1**d**), as previously described ^40^. The resulting nucleosomes were tested for their interaction with purified MutSβ and MutSα. Although MutSβ was pulled down by both the H3K56-nucleosome and H3K56ac-nucleosome, the amount of MSH3 pulled down by the H3K56-nucleosome is 4-fold more than that pulled down by the H3K56ac-nucleosome (Figure 1**e**, compare lane 1 with lane 2). This pulldown appears to be MutSβ-specific, as no MSH6 (MutSα) was precipitated in either nucleosome reaction (Figure 1**e**, lanes 3 and 4). Taken together, these data presented here strongly suggest that MutSβ interacts with chromatin through unmodified H3K56, but this interaction is inhibited when H3K56 is acetylated.

### H3K56 deacetylation promotes MutSβ chromatin binding

To determine the impact of H3K56 acetylation and deacetylation on MutSβ’s chromatin recruitment, we treated HeLa cells with histone deacetylase inhibitor Romidepsin (FK228), and measured MSH3 chromatin localization using immunofluorescence assays. In comparison to untreated cells, FK228-treated cells exhibited increased levels of H3K56ac; the FK228-induced increase in the H3K56ac abundance almost completely deprived MSH3 from chromatin binding (Figure 2**a**). It is interesting to note that even in control cells, a high level of H3K56ac is accompanied with a low level of chromatin-bound MSH3. The reverse is also true, i.e., high levels of MSH3 are coupled with low levels of H3K56ac (Figure 2**a**, upper). This mutually exclusive relationship between H3K56ac and MSH3 was confirmed when measuring the amount of MSH3 and H3K56ac in chromatin fraction by Western blotting (Figure 2**b**).

**Fig. 2.**
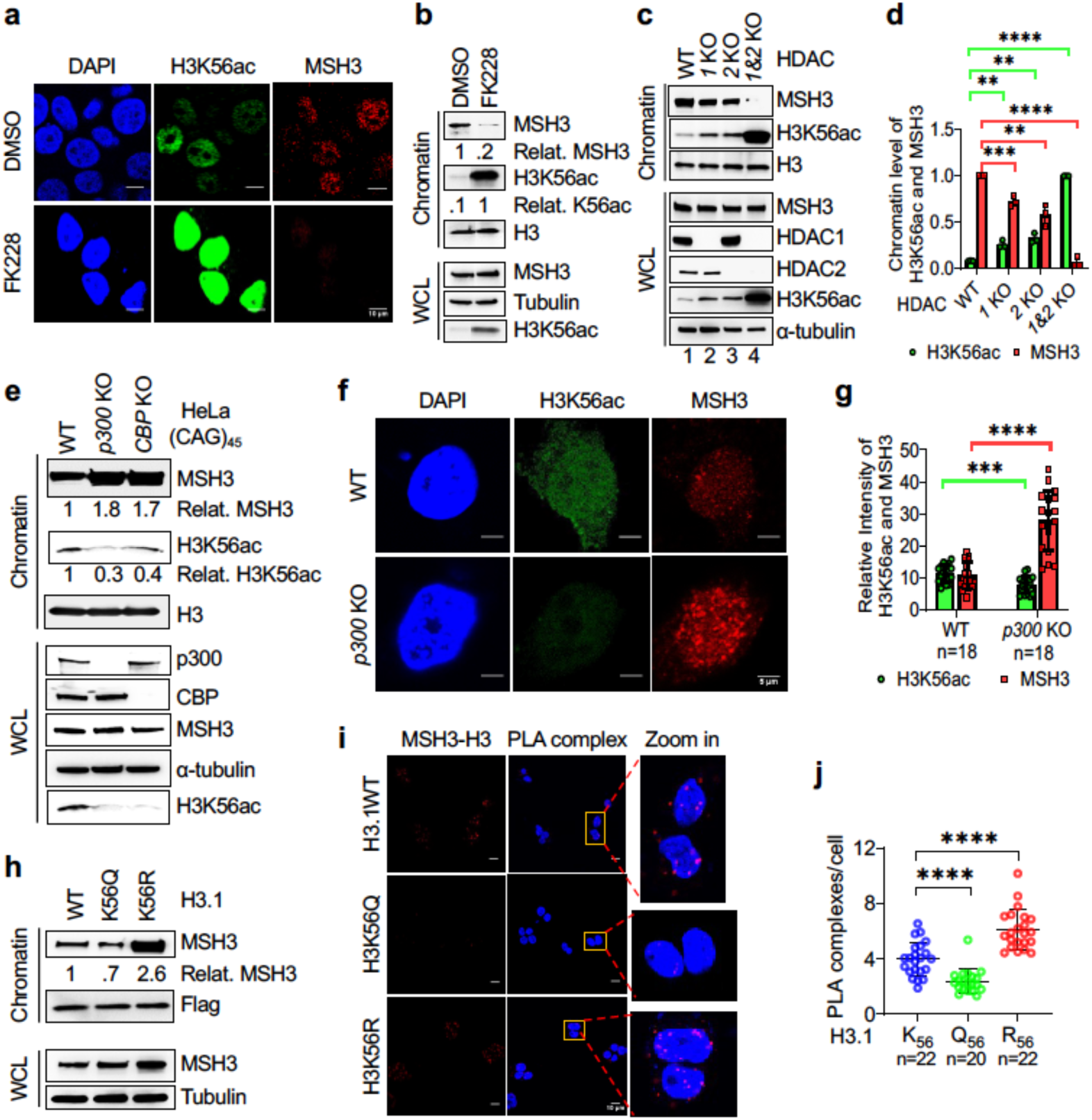
H3K56 deacetylation facilitates MutSβ chromatin recruitment. **a,** Representative immunofluorescence images showing inverse chromatin intensity between MSH3 and H3K56ac in HeLa cells treated with or without HDAC1/2 inhibitor FK228. **b,** Western blots showing inverse correlation between the abundance of H3K56ac and MSH3 in chromatin fraction. HeLa cells were treated with or without 0.2 mM HDAC inhibitor FK228 or the same volume of DMSO for 24 hours, and cell lysates and chromatin fractions were analyzed for MSH3 and H3K56ac by Western blotting analysis. **c,** Western blots showing H3K56ac abundance and chromatin-bound MSH3 in HeLa WT (WT), *HDAC1* KO (1 KO), *HDAC2* KO (2KO) and *HDAC1-HDAC2* double KO (1&2 KO). **d,** Quantification of H3K56ac and chromatin-bound MSH3 levels, as shown in C. Data are shown as mean ± SD. ** *P<* 0.01, *** *P<* 0.001, and **** *P* <0.0001 were determined by unpaired t-test. **e,** Western blots showing enhanced chromatin-bound MSH3, but reduced level of H3K56ac in HeLa cells with *p300* KO. **f,** Representative Immunofluorescence images showing reduced H3K56ac abundance and increased level of MSH3 foci in HeLa *p300* KO cells, as compared with WT cells. **g,** Quantification results showing relative H3K56ac and chromatin-bound MSH3 levels, as shown in F. Data are shown as mean ± SD. *** *P<* 0.001, and **** *P* <0.0001 were determined by unpaired t-test. **h,** Western blotting analysis to show the levels of chromatin-bound MSH3 in HEK293 cells stably expressing H3.1 WT, acetylation-mimicking H3.1-K56Q or deacetylation-mimicking H3.1-K56R. **i,** Representative PLA images to show specific interaction between MSH3 and ectopic H3 in HEK293 cells expressing H3.1 WT and H3K56R, but not in HEK293 cells expressing H3K56Q. **j,** Quantification of the number of PLA complexes in HEK293 cells expressing 3.1WT, H3K56Q and H3K56R, as indicated. Unpaired t-test was used in statistical analysis. Data were from three independent experiments, and shown as mean ± SD. ** *P<* 0.001, *** *P<* 0.001, and **** *P* <0.0001 were determined by unpaired t-test.

To further determine the effect of H3K56 acetylation status on MutSβ’s chromatin binding, we knocked out histone deacetylases HDAC1, HDAC2, or both, which have been shown to remove the acetyl group from H3K56ac ^41,42^. We analyzed MSH3 chromatin association in these knockout HeLa cells. Although HDAC1 knockout (1 KO) or HDAC2 knockout (2 KO) enhanced H3K56ac levels in both whole cell lysates and chromatin, the HDAC1-HDAC2 double knockout (1&2 KO) significantly increased the H3K56ac level (Figures 2**c** and 2**d**); and this increase significantly decreased the chromatin-bound level of MSH3, as essentially no MSH3 was recovered in the chromatin fraction derived from the double knockout cells (Figure 2**c**, lane 4). We next separately knocked out histone acetyltransferases p300 and CBP, both of which acetylate H3K56 ^43,44^, as we failed to obtain a *p300-BCP* double knockout clone. Consistent with a previous study ^43^, either knockout resulted in dramatical reduction in H3K56ac levels, but significant increase in MSH3 chromatin levels (Figures 2**e**, 2**f** and 2**g**). Therefore, H3K56 acetylation status controls MutSβ’s chromatin localization.

To orthogonally test this notion, we generated HEK293 cell lines stably expressing Flag-tagged histone H3.1 proteins that carry wild type K56 (WT) or a substitution of K56 to glutamine (K56Q) or arginine (K56R) (Figure S2**a**). The K56Q substitution mimics a constitutively acetylated K56, while the K56R substitution mimics a constitutively unmodified K56 ^45–47^. We found that the amount of the chromatin-bound MSH3 in cells expressing H3.1-K56R was ∼3.6-fold higher than that in cells expressing H3.1-K56Q (Figure 2**h**). In situ proximity ligation assay (PLA), which detects protein-protein interactions with high specificity and sensitivity ^48^, revealed that cells expressing the acetylation-mimicking H3.1-K56Q mutant protein showed significantly decreased level of PLA complexes, as compared with cells expressing WT H3.1 and the deacetylation-mimicking H3.1-K56R, the latter of which showed the highest level of PLA complexes (Figures 2**i** and **j**). Collectively, the results shown here strongly suggest that H3K56 acetylation inhibits, but its deacetylation promotes MutSβ chromatin binding *in vivo*.

### MSH3 domain II is required for MutSβ binding to chromatin

To identify the MSH3 domain that interacts with H3K56, we generated seven MSH3 fragments (Figure 3**a**, F1-F7) according to the known structural domains ^20^ and expressed them as GST fusion proteins. The resulting fusion proteins were examined for interacting with the biotinylated K56(44-63) peptide in GST-pulldown assays. We found that only fragments F1 and F4 efficiently pulled down the K56(44-63) peptide (Figure 3**b**). Since F1 and F4 share MSH3 residues 343-531 (domain II), the results suggest that MSH3 residues 343-531 are involved in the MutSβ-H3K56 interaction. If this is true, MutSβ depleted of MSH3 domain II (i.e., F4) will not be recruited to chromatin.

**Fig. 3.**
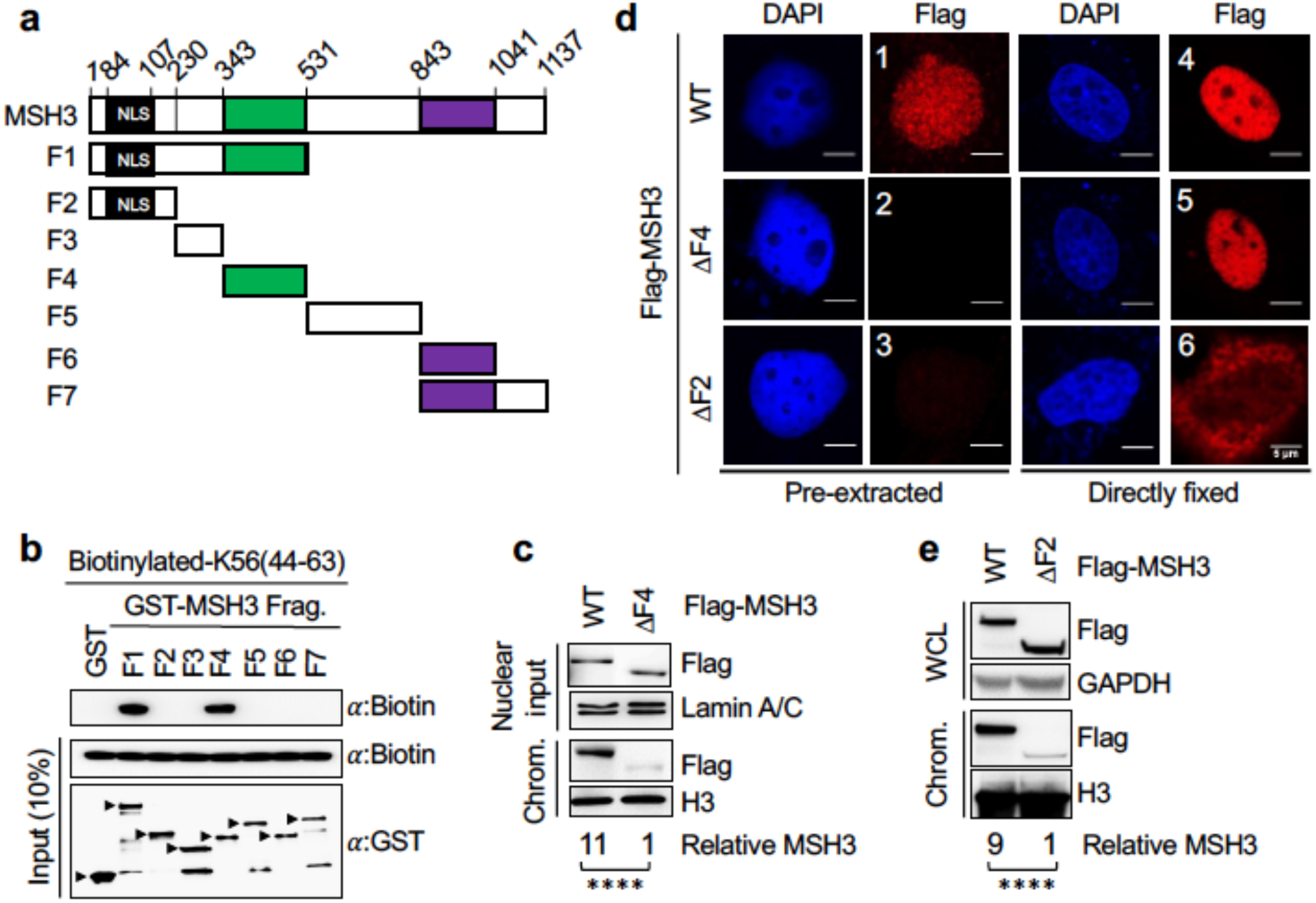
MutSβ interacts with chromatin via MSH3 residues 344-531. **a,** Schematic diagram of the GST-tagged MSH3 fragments used in the GST pulldown assay. **b,** GST pulldown-Western blotting analysis showing specific interaction between H3K56 peptide and GST-MSH3 fragments containing fragment 4 (residues 344-531). The pulldown H3 peptide was detected by an antibody against biotin. **c,** Western blots showing that MSH3 residues 344-531 are required for MSH3 chromatin binding. Flag-MSH3 or Flag-ΔF4-MSH3 was expressed in HeLa-*MSH3* KO cells. Nuclear and chromatin fractions were analyzed for MSH3 by Western blotting analysis, as indicated. ****, p<0.001. **d,** Representative immunofluorescence images showing subcellular compartment localization of Flag-MSH3, Flag-ΔF4-MSH3, and Flag-ΔF2-MSH3 in HeLa-*MSH3* KO cells with (left) or without (right) the pre-extraction by the CSK buffer, which removes soluble proteins. **e,** Western blot showing the expression of Flag-MSH3 and Flag-ΔF2-MSH3 in cell extract and chromatin fractions. Flag-MSH3 or Flag-ΔF2-MSH3 was expressed in HeLa-*MSH3* KO cells. ****, p<0.001.

To test this idea, we expressed Flag-tagged full length MSH3 (WT) and F4-deleted MSH3 (ΔF4-MSH3) in *MSH3*-knockout (MSH3 KO) HeLa cells and determined their chromatin binding activity. Western blotting analysis showed that although roughly equal amount of MSH3 and ΔF4-MSH3 were present in the nucleoplasm, the level of the chromatin-bound MSH3 is significantly higher than that of the chromatin-bound ΔF4-MSH3 (Figure 3**c**). We then determined chromatin-bound MSH3 by confocal immunofluorescence analysis after cells were pre-extracted with the buffer that removes loosely chromatin-bound proteins before fixing ^49,50^. The results revealed that WT MSH3 remained chromatin bound (Figure 3**d**, image 1), but little ΔF4-MSH3 was recovered in the nucleus (Figure 3**d**, image 2), suggesting that MutSβ with a ΔF4-MSH3 subunit has lost its chromatin binding activity.

To make sure that deleting F4 did not affect ΔF4-MSH3 to enter into the nucleus, we generated a fragment 2-depleted MSH3 (ΔF2-MSH3), which lacks a major nuclear localization sequence (NLS) (see Figure 3**a**) and thereby is unable to enter into the nucleus. Like Flag-tagged WT and ΔF4-MSH3, Flag-tagged ΔF2-MSH3 was well expressed in HeLa-MSH3 KO cells (Figure 3**e**). As expected, WT MSH3 was detected in the nucleus (Figure 3**d**, image 4); ΔF2-MSH3 was found in cytoplasm (Figure 3**d**, image 6), but was completely removed from cytoplasm upon pre-extraction (Figure 3**d**, image 3). However, like WT MSH3, ΔF4-MSH3 was also seen in the nucleus (Figure 3**d**, image 5), indicating that ΔF4-MSH3 can effectively translocate into the nucleus. Therefore, we conclude that MutSβ interacts with chromatin through its MSH3 domain II or residues 343-531.

### An MSH3 aromatic packet is responsible for interacting with H3K56

Disordered regions often play important roles in the function of chromatin-associated proteins, and many of these regions, particularly their nearby aromatic residues, can form a packet or channel to mediate protein-chromatin interactions ^51,52^. This has been exemplified in the interactions between the MSH6-PWWP domain and H3K36me3 ^23^, and the histone acetyltransferase Rtt109 and H3K56 ^53^. In the latter case, Rtt109 residues Y145, R140 and F138 form an aromatic packet surrounding the aliphatic portion of H3K56 (Figure 4**a**) and facilitate acetylation of K56 ^53^. To determine whether a similar mechanism is involved in the interaction between MSH3 and H3K56, we analyzed the structure of the MSH3 domain II ^20^ and identified two disordered regions, which are solvent exposed and not involved in either MSH2 or mismatched DNA substrate binding (Figure 4**b**). Region 1 consists of amino acid residues 367-375 (ENVRDKKKG), and is surrounded by aromatic residues F378, Y456 and F460. Region 2 is composed of residues 469-482 (KDTVDIKGSQIISG), which is adjacent to aromatic residues F466, Y467 and F393 (Figure 4**b**). Based on the structure of the MSH3 domain II and the Rtt109-H3K56 interaction, we simulated an MSH3-H3K56 interaction model, where aromatic residues F378, F393, F516, F466, Y456 and F460 of MSH3 form a binding channel that accommodates H3 residues 54-59, with MSH3 F393 and F516 functioning as a supporting role (Figure 4**c**).

**Fig. 4.**
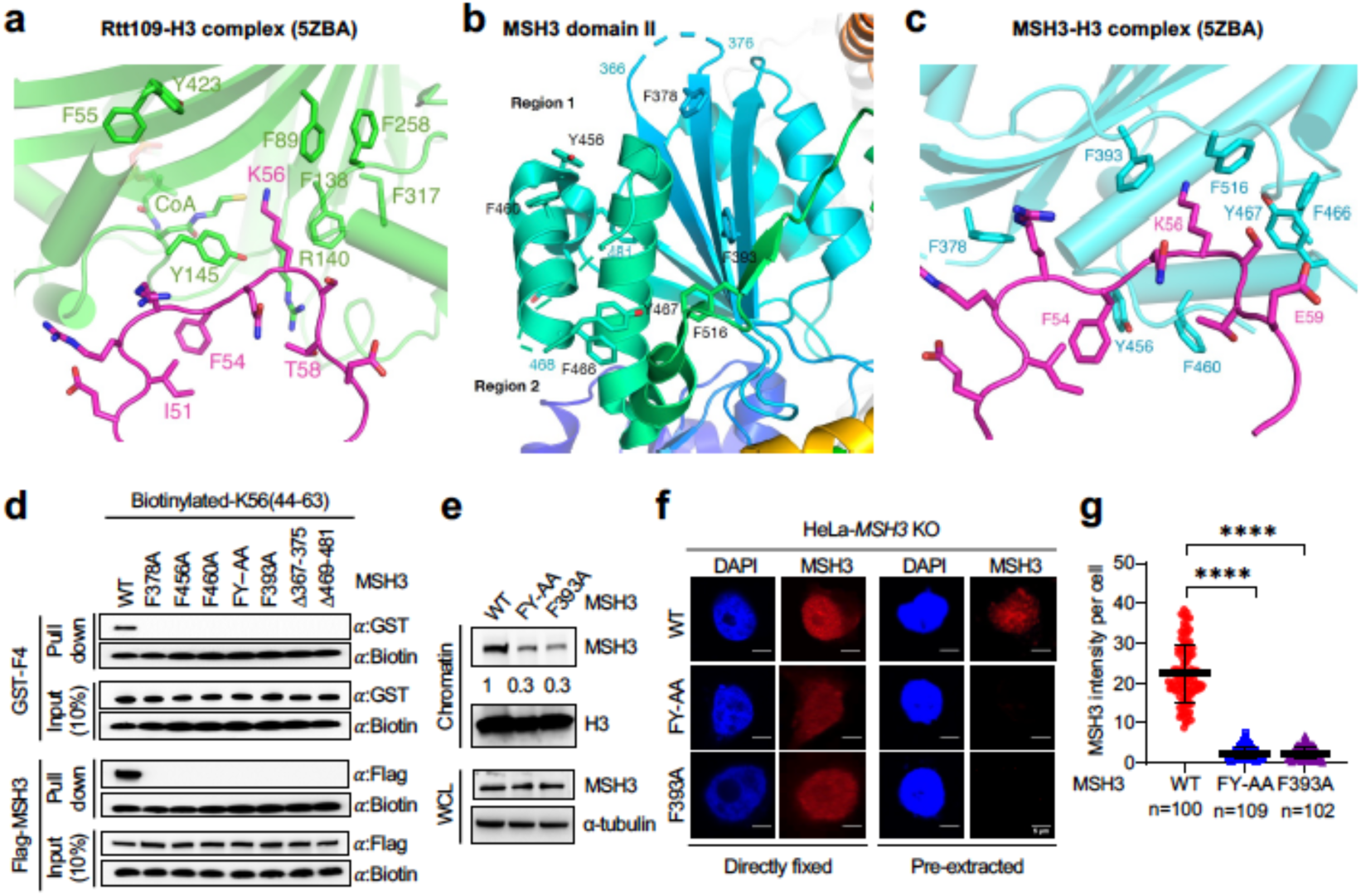
The H3K56-MutSβ interaction relies on the aromatic packet of MSH3 domain II. **a,** The interactions between Rtt109 and H3K56 according to the crystal structure of Rtt109-Asf1-H3-H4-CoA complex (PDB: 5ZBA). Rtt109 is shown in light green and H3 in magenta cartoons. Critical residues at the interface are highlighted in stick model and labeled. **b**, The structure of MSH3 domain II (PDB: 3THX) is shown as cartoon in blue to green color gradient indicating the N to C-terminal sequence. The disordered regions 1 and 2 are indicated by the dashed lines, and surrounding aromatic residues are shown in stick model and labeled. **c,** A complex between MSH3 domain II and the H3K56 peptide is simulated based on the Rtt109-H3 structure. Both disordered regions are involved. F383, and F516 sandwich K56, and F516 is supported by F466 and Y467. **d,** Biotin-streptavidin pulldown assay showing the requirement for a functional MSH3 aromatic packet in the H3K56-MutSβ interaction. Biotinylated-H3K56(44-63) peptide was incubated with various isoforms of GST-tagged MSH3 fragment 4 (GST-F4) (upper) or individual derivatives of Flag-MSH3 (bottom), and the pulldown species i.e., GST-F4 or Flag-MSH3 derivatives, were analyzed by Western blotting and detected by a GST or Flag antibody, respectively. **e,** Western blots showing the chromatin binding levels of Flag-MSH3 and its aromatic packet mutants. **f,** Representative immunofluorescence images showing the chromatin binding activity of Flag-MSH3 and its aromatic packet mutants HeLa-*MSH3* KO cells, with (right) or without (left) pre-extraction by the CSK buffer before fixing. **g,** Quantification of the MSH3 chromatin intensity of Flag-MSH3 and its aromatic packet mutants in HeLa-*MSH3* KO cells with the CSK buffer pre-extraction. Data are shown as mean ± SD. ****, *P* <0.0001.

To test this model, we created several MSH3 mutant proteins (both full length and F4) that carry either a deletion of a disordered region (i.e., MSH3 residues 367-375 or 469-481) or an alanine substitution of aromatic residues F378, F456, F460, F466, Y467 or F393 (Figure S2**b**-**h**). The full length MSH3 proteins were Flag-tagged, while the MSH3 F4 isoforms were GST-fused. We first examined the individual MSH3 proteins for their ability to interact with biotinylated H3 peptide K56(44-63) using the biotin-streptavidin pulldown assay, as described above.

Interestingly, we found that although WT GST-MSH3-F4 and WT Flag-MSH3 were efficiently pulled down by the H3 peptide K56(44-63), all mutants, regardless of the depletion of a disordered region or substitution of an aromatic residue, failed to interact with the H3 peptide (Figure 4**d**). Consistently, alanine substitutions of aromatic residues F393 or F466 and Y467 significantly reduced the MutSβ’s chromatin binding activity (Figure 4**e**). Confocal immunofluorescence analysis showed that although both WT and mutant MSH3s entered the nucleus efficiently (Figure 4**f**, left), only WT MSH3 was found to tightly bind to chromatin (Figure 4**f**, right; and Figure 4**g**). These results suggest that the aromatic channel formed by both the disordered regions and their surrounding aromatic residues are essential for the MutSβ-H3K56 interaction.

### Disrupting the MutSβ-H3K56 interaction stabilizes CAG repeats

Given MutSβ’s role in promoting TNR instability, we propose that blocking the MutSβ chromatin recruitment would stabilize (CAG)•(CTG) repeats. To test this hypothesis, we established stable expression of WT MSH3, F4-deleted MSH3 (ΔF4) and F_466_Y_467_–AA-MSH3 (FY-AA) in MSH3-knockout HeLa cells with 45 (CAG)•(CTG) repeats alongside a single ectopic copy of the c-Myc replication origin ^54^ (Figure 5**a**). The resulting individual clones were cultured for 100 passages, and genomic DNAs isolated from the individual clones were used to analyze the (CAG)_45_ stability by PCR. Both contracted and expended CAG repeats were observed in HeLa-(CAG)_45_ clones expressing WT MSH3, which is consistent with previous studies ^54^; however, the contracted and expanded CAG repeat species were not detected in cells expressing either FY-AA or ΔF4 MSH3 (Figure 5**b**). These results confirm that disrupting the MutSβ-H3K56 interaction indeed facilitates the TNR stability.

**Fig. 5.**
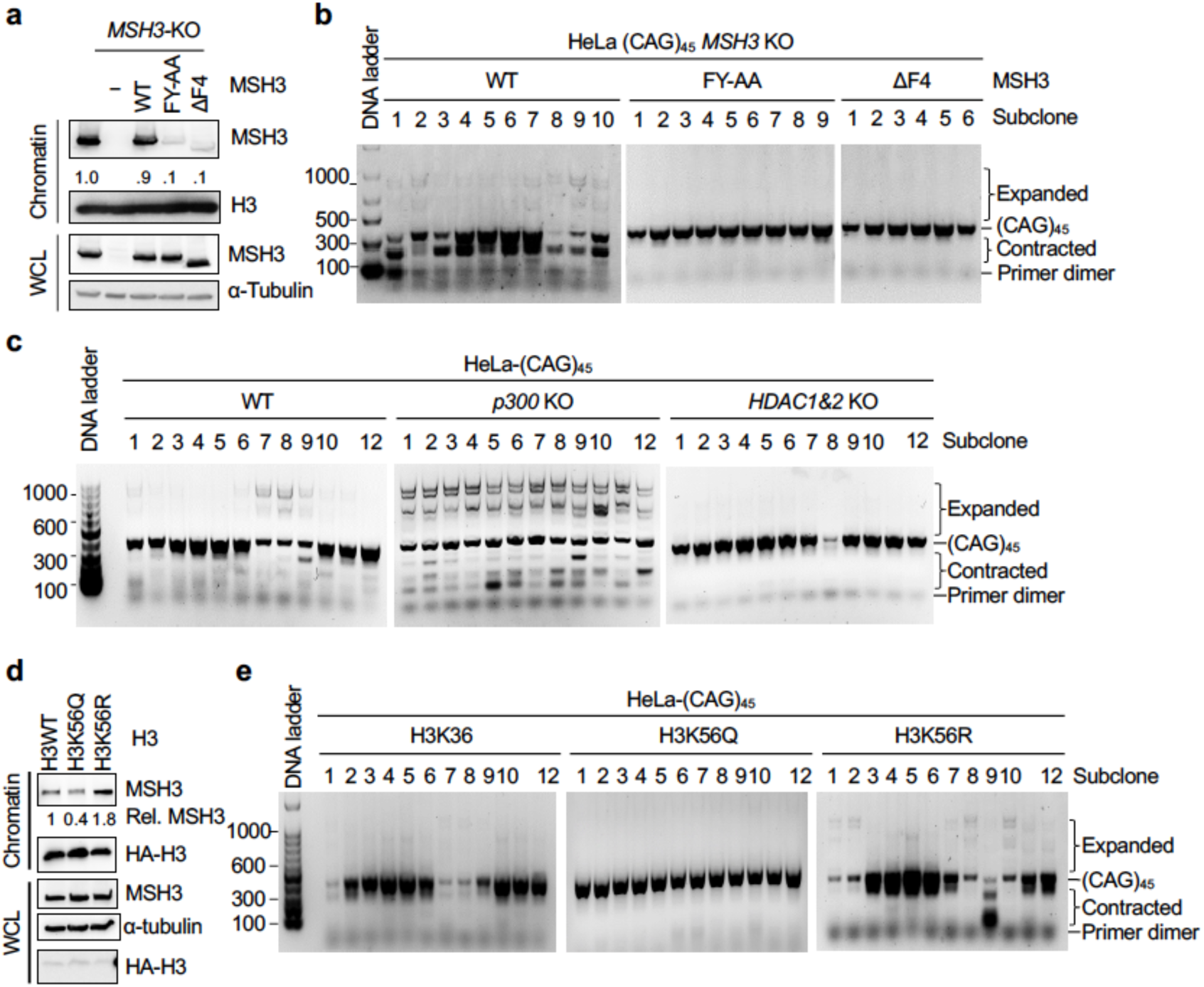
The MutSβ-H3K56 interaction promotes (CAG)•(CTG) repeat instability. **a,** Western blots showing stable expression of MSH3 or its aromatic packet mutants in HeLa-(CAG)_45_-*MSH3* KO cells. **b,** PCR analysis to detect the (CAG)_45_ stability in sub-clones of HeLa-(CAG)_45_ *MSH3* KO cells stably expressing MSH3 (WT), F4-deleted MSH3 (ΔF4) or F_466_Y_467_–AA-MSH3 (FY-AA). Genome DNAs isolated from individual clones after 100 passages were used to amplify the CAG repeat sequence, as described previously ^54^. PCR product was resolved in 1.5% agarose gel and detected by GelGreen staining. **c,** PCR analysis showing the (CAG)_45_ stability in sub-clones of HeLa-(CAG)_45_ cells depleted of *p300* (*p300* KO) or *HDAC*1&2 (*HDAC*1&2 KO). Cells were cultured for 160 passages before harvest. PCR analysis was performed as described in B. **d,** Western blots displaying the levels of chromatin-bound MSH3 in HeLa-(CAG)_45_-*MSH3* KO cells stably expressing H3 WT, acetylation-mimicking H3K56Q or deacetylation-mimicking H3K56R. **e,** PCR analysis showing the (CAG)_45_ stability in sub-clones of HeLa-(CAG)_45_ cells stably expressing H3K56, H3K56Q and H3K56R, as indicated. Cells were cultured for 140 passages before harvest. PCR analysis was performed as described in B.

### Histone deacetylation promotes CAG repeat instability

Since the acetylation status of H3K56 regulates the MutSβ-H3K56 interaction (Figure 2), we speculated that histone deacetylation or acetylation promotes (CAG)•(CTG) repeat instability or stability, respectively. To explore these possibilities, we analyzed the (CAG)_45_ sequence stability in HeLa-(CAG)_45_ cell clones with *p300* knockout (*p300* KO) or *HDAC1&2* double knockout (*HDAC1&2* KO). The former cells (*p300* KO) showed lower levels of H3K56ac, but higher levels of chromatin-bound MSH3 (Figure 2**f**), while the latter cells (*HDAC1&2* KO) displayed higher levels of H3K56ac, but lower levels of chromatin-bound MSH3 (Figure 2**d**). In comparison with WT clones, some of which showed limited expansion or contraction of CAG repeats (Figure 5**c**, left gel), all *p300* KO clones analyzed exhibited elevated levels of repeat expansions, as well as contractions (Figure 5**c**, middle gel); however, repeat expansions and contractions were not detectable in *HDAC1&2* KO cells (Figure 5**c**, right gel).

To delve deeper into the influence of H3K56’s status on CAG repeat instability, we established HeLa-(CAG)45 cells stably expressing a HA-tagged histone H3.3 protein carrying wild-type K56 (WT), acetylation-mimicking K56Q mutation or non-acetylation mimicking K56R mutation (Figure 5**d**), and analyzed the stability of the (CAG)45 sequence. As illustrated in Figure 5**e**, cells carrying H3K56R (right gel) exhibited significantly increased CAG repeat instability compared to cells carrying K56 (left gel) or K56Q (middle gel).

Taken together, these results suggest that histone deacetylation, which facilitates MutSβ chromatin binding via interacting with H3K56 (Figure 2), destabilizes CAG repeats, while histone acetylation, which blocks MutSβ’s interaction with H3K56, stabilizes CAG repeats.

### The MutSβ-H3K56 interaction facilitates expansion of *HTT* CAG repeats and mHtt aggregation

To determine the impact of the MutSβ-H3K56 interaction on expansion of CAG repeats in the *HTT* gene, which causes HD ^55,56^, we utilized two striatal cell lines, STHdh^Q7/Q7^ and STHdh^Q111/Q111^, which were derived from a knock-in transgenic mouse containing homozygous *HTT* loci with humanized exon 1 carrying 7 and 111 CAG repeats, respectively. Given the conservation of Lys 56 in histone 3 between human and mouse (Figure S3**a**), we established STHdh^Q7/Q7^ and STHdh^Q111/Q111^ cell lines stably expressing HA-tagged histone H3.3 that carries K56, K56Q or K56R. Similar to human cells, the chromatin-bound Msh3 is significantly elevated in both STHdh^Q7/Q7^ and STHdh^Q111/Q111^ cells expressing the non-acetylation mimicking K56R, compared to cells expressing WT K56 or the acetylation-mimicking K56Q (Figure 6**a**). We then investigated the impact of the differential MutSβ recruitment on the instability of *HTT* CAG repeats in STHdh^Q7/Q7^ and STHdh^Q111/Q111^ cells. PCR analysis targeting the CAG repeats in humanized *HTT* exon 1 revealed that STHdh^Q7/Q7^ and STHdh^Q111/Q111^ cells harboring K36, K36Q and K56R displayed distinct differences in *HTT* CAG repeat numbers (Figure 6**b**). Consistent with what was observed in human cells (Figures 5**e**), both STHdh^Q7/Q7^ and STHdh^Q111/Q111^ cells with H3K36 also showed a mild repeat instability, especially in STHdh^Q7/Q7^ demonstrating a maximum CAG repeat number of 11 (from Q7 to Q11) after 12 weeks of culturing; however, over the same duration of cell culturing, cells harboring H3K56Q mutation displayed no expanded CAG repeats (in STHdh^Q7/Q7^ cells) or very limitedly expanded CAG repeats (in STHdh^Q111/Q111^); but those expressing deacetylation mimic H3K56R displayed multiple expanded CAG repeats, reaching a maximum number of 18 in STHdh^Q7/Q7^ cultured for 12 weeks and the maximum number of 144 in STHdh^Q111/Q111^ cells cultured for 22 weeks (Figure 6**b**). We further quantified the level of repeat instability in STHdh^Q111/Q111^ cells using the Gene-Mapper Software ^57^, which tracks all peak heights and eliminates peaks whose heights are below the 20% threshold (see red dosh line) from quantification. The results demonstrated a significantly higher level of the instability index in STHdh^Q111/Q111^ cells expressing H3K56R, compared to those expressing WT H3K56 and H3K56Q (Figure 6**c**).

**Fig. 6.**
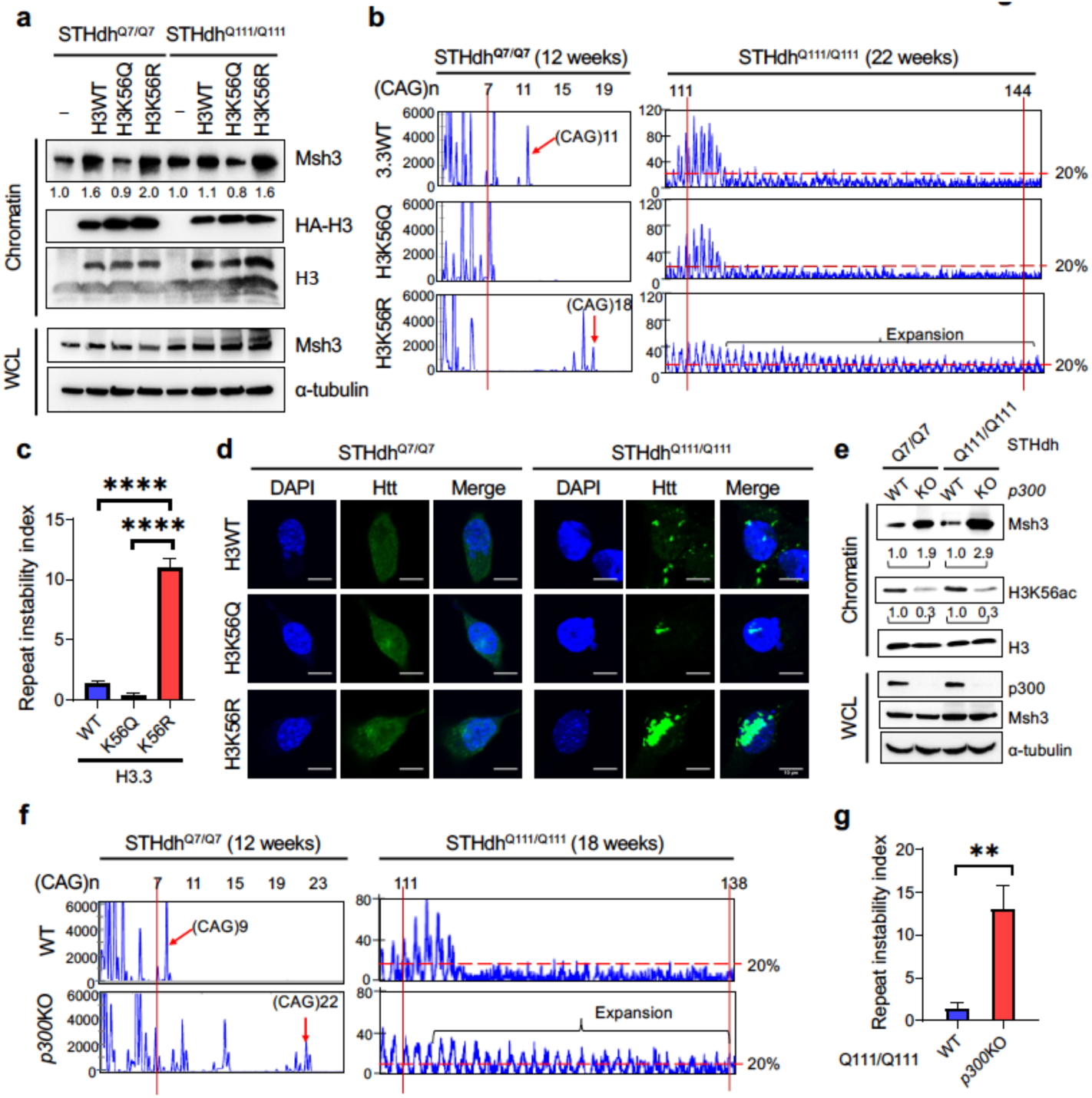
MutSβ chromatin recruitment induces expansion of *HTT* (CAG)_n_ and mHtt aggregation. **a,** Western blots displaying the expression of Msh3 in chromatin fractions and whole cell lysates (WCL) in STHdh^Q7/Q7^ and STHdh^Q111/Q111^ cells. WT and stable cell lines expressing HA-tagged histone H3K56, H3K56Q and H3R56 were indicated. **b,** Representative Gene Mapper profiles illustrating *HTT* CAG repeat size distributions in STHdh^Q7/Q7^ and STHdh^Q111/Q111^ cells stably expressing H3K56, H3K56Q, and H3K56R. STHdh^Q7/Q7^ cells were cultured over 12 weeks for *HTT* CAG repeat analysis (left panel), while STHdh^Q111/Q111^ cells were cultured over 22 weeks for *HTT* CAG repeat analysis (right panel). **c,** Quantification of CAG repeat instability index in STHdh^Q111/Q111^ cells stably expressing H3K56, H3K56Q, and H3K56R. **d,** Representative immunofluorescence images showing Htt staining in STHdh^Q7/Q7^ (left panel) and STHdh^Q111/Q111^ (right panel) cells stably expressing H3K56, H3K56Q, and H3K56R. **e,** Western blots displaying the expression of Msh3 in chromatin fractions and whole cell lysates (WCL) in STHdh^Q7/Q7^ and STHdh^Q111/Q111^ cells. WT and *p300*-KO cell lines were indicated. **f,** Representative Gene Mapper profiles illustrating *HTT* CAG repeat size distributions in WT and *p300*-KO cell lines with Q7/Q7 (left panel) or Q111/Q111 (right panel). **g,** Quantification of CAG repeat instability index in WT and *p300*-KO STHdh^Q111/Q111^ cells. Mean ± SD values were shown. **, p<0.01; ****, p<0.0001.

The CAG repeats in the *HTT* gene are translated into a polyglutamine (polyQ) stretch in the N-terminal region of the huntingtin (Htt) protein. In Huntington’s disease (HD) patients, the number of CAG repeats is expanded to ≥36, resulting in the production of a mutated Htt (mHtt) protein with an expanded polyQ region ^58^. The rate of mHtt protein aggregation correlates with the length of the polyQ expansion ^59,60^, making the aggregation of mHtt a hallmark feature of HD ^61,62^. Since the recruitment of MutSβ promotes CAG repeat expansion in the *HTT* gene, we hypothesized that this mechanism would result in an expansion of the polyQ lengths within mHtt, consequently exacerbating mHtt aggregation. To test this hypothesis, we examined Htt aggregation in STHdh^Q7/Q7^ and STHdh^Q111/Q111^ cells. As shown in Figure 6**d**, there is no evident Htt aggregation in STHdh^Q7/Q7^ cells. Although STHdh^Q7/Q7^ cells harboring H3K56R at 12 weeks displayed an apparent CAG repeat expansion, the maximum of CAG repeat length was only 18, which remains insufficient to trigger Htt aggregation. This result is consistent with previous study showing that Htt protein aggregation typically manifests when the polyQ tract exceeds approximately 40 glutamine residues ^63^. As anticipated, STHdh^Q111/Q111^ cells exhibited mHtt aggregation. Remarkably, STHdh^Q111/Q111^ cells carrying the H3K56R mutation displayed extensive mHtt aggregation, including the formation of inclusion bodies (Figure 6**d**). This result aligns with prior studies demonstrating that the length of the polyQ tract influences the extent of mHtt aggregation ^60^ and the accumulation of mHtt within intracellular inclusion bodies ^64^.

Similarly to human cells, the chromatin-bound Msh3 was notably elevated in both STHdh^Q7/Q7^and STHdh^Q111/Q111^ cells when the histone acetyltransferase p300 was depleted, compared to WT cells (Figure 6**e**). Correspondingly, we observed that both STHdh^Q7/Q7^and STHdh^Q111/Q111^ cells exhibited significantly expanded CAG repeats when *p300* was depleted, in comparison to WT cells with the same duration of cell culturing (Figure 6**f**). STHdh^Q111/Q111^ cells with p300 knockout showed significantly higher CAG repeat instability index, as compared with their WT controls (Figure 6**g**). We further showed that STHdh^Q111/Q111^cells, upon depletion of p300, exhibited extensive mHtt aggregation compared to wild-type STHdh^Q111/Q111^ cells after 18 weeks of cell culturing (Figure S3**b**).

Taken together, these results further suggest that H3K56 acetylation status controls the chromatin recruitment of the CAG repeat instability driver, MutSβ, with acetylation inhibiting but deacetylation promoting the recruitment, and that localization of MutSβ by H3K56 stimulates the expansion of CAG repeats in the *HTT* gene and mHtt aggregation.

## Discussion

MutSβ is known to promote TNR instability, particularly the expansion of CAG repeats in the *HTT* gene by directly interacting with DNA. To execute this function, MutSβ must be located on chromatin so that it has an easy access to DNA. However, the molecular mechanism by which MutSβ is recruited to chromatin is unknown until this study, where H3K56 is identified as the histone mark engaging MutSβ to chromatin through its physical interaction with an aromatic packet in the MSH3 subunit of MutSβ. The binding of MutSβ to chromatin is regulated by the acetylation status of H3K56, with acetylated H3K56 inhibiting but deacetylated H3K56 promoting MutSβ chromatin recruitment. We also show that the H3K56-MutSβ interaction directly controls the instability of CAG repeats, including the expansion of the very short *HTT* (CAG)7, as blocking the interaction by disrupting the MutSβ’s aromatic packet or acetylating H3K56 stabilizes CAG repeats, while deacetylating H3K56 stimulates CAG repeat instability.

The identification of H3K56 as the histone mark for MutSβ’s chromatin engagement indicates that MutSα and MutSβ are independently recruited to chromatin. This is very important for genome maintenance, because MutSα and MutSβ are required to correct base-base mismatches and small insertion-deletion mispairs generated during DNA replication, respectively ^21,65^. Whereas MutSα relies on the MSH6 PWWP domain for chromatin recruitment ^23^, the MutSβ chromatin engagement requires the MSH3 aromatic packet. Mutations in the PWWP domain block the H3K36me3-MutSα interaction and lead to a mutator phenotype ^23–25^.

Similarly, we show here that disrupting the MSH3 aromatic packet also interrupts the H3K56-MutSβ interaction. Like the MSH6 PWWP domain, which is not present in MSH2 and MSH3, the aromatic packet, which consists of two disordered regions (Supplemental Figure 1**a**, red-type) and several critical aromatic residues (Figure S1**a**, blue type), exists in MSH3, but not in MSH2 and MSH6 (Figure S1**a**). Consistently, the result from the GST pulldown assay showed that the MSH6 domain II failed to interact with H3K56(44-63) peptide (Figure S1**b**). However, unlike the critical genome-caretaker MutSα, MutSβ can promote TNR instability, particularly when MSH3 is overexpressed. This is probably the reason why human cells express limited amounts of MSH3 ^66,67^ to effectively control its TNR expansion function.

The identification of the H3K56-MutSβ interaction opens new approaches against HD and other TNR expansion-caused diseases. Given that MutSβ’s ATPase activity is crucial for CAG repeat expansions ^68^ and that MutSβ’s cohesive interactions with the downstream MMR factors such as MutL homologs and exonuclease 1 are important for MMR, the current HD therapeutical efforts have been vigorously focused on identifying small molecules to inhibit MutSβ’s enzymatic activities ^17,18^ (also see https://chdifoundation.org/dna-repair-handling/). Unfortunately, even if these interventions could efficiently target MutSβ, they are likely inhibiting the same activities of MutSα. This is because MutSα and MutSβ adopt essentially the same ATP binding/hydrolysis structure ^19,20^ and use the same manner to interact with their downstream factors during MMR ^21,22^. Since defects in MSH6, but not MSH3, cause a mutator phenotype and cancer development ^6–9^, MutSβ’s enzymatic inhibitors, which also inhibit MutSα’s activities, are expected to be very detrimental to patients. However, our finding here provides opportunities to develop MutSβ-specific inhibitors, i.e., by blocking the H3K56-MutSβ interaction.

HDAC inhibitors have been shown to suppress CAG repeat expansion ^27–30^, but the molecular basis is unknown. Our study has solved this puzzle. This is because HDAC inhibitors enhance H3K56 acetylation levels, which blocks the H3K56-MutSβ interaction, thereby eliminating MutSβ from chromatin binding (Figures 1 and 2). However, it is well documented that HDAC inhibitors alter not only acetylation of H3K56, but also acetylation of other lysine residues of histones, as well as many other proteins involved in regulating gene expression, cell proliferation and activation of cell death via apoptosis and autophagy ^35,69^. Thus, although HDAC inhibitors can inhibit CAG repeat expansions, they likely cause severe side effects. Based on the fact that disruption of the disordered regions or substitution of any aromatic residues in the MSH3 domain II interrupts the H3K56-MutSβ interaction, which promotes CAG repeat stability (Figure 5), a second but side effect-free way to block the H3K56-MutSβ interaction is to identify small molecules to alter the structure of the MSH3 aromatic packet. Therefore, this study provides an innovative approach to develop drugs as an effective therapy for HD and TNR expansion diseases by specifically inhibiting MutSβ repeat expansion function.

## Materials and Methods

### Preparation of MSH3 and MSH6 constructs

The full-length *MSH3* gene, fragment 2-deleted *MSH3*, fragment 4-deleted *MSH3*, and the individual aromatic mutants of full-length *MSH3* were generated by PCR. The PCR products were cloned into pCMV5-Flag vector to express Flag-tagged MSH3 derivatives in HeLa *MSH3* knockout cells. MSH6 domain II, MSH3 fragment 4 wild-type and aromatic mutants were similarly generated by PCR, and PCR products were cloned into pGEX-4T-2 vector for express GST-tagged proteins in *Escherichia coli* strain BL21.

### Cell lines and cell culture

HeLa cell lines used in this study are HeLa carrying 45 CAG repeats in the c-myc replication origin (HeLa-(CAG)_45_). HeLa(CAG)_45_ and HEK293 cells were cultured in Dulbecco’s modified Eagle’s medium (DMEM) supplemented with 10% fetal bovine serum (FBS) and 4 mM L-glutamine at 37 °C in a humidified atmosphere with 5% CO_2_.

STHdh^Q^^7/Q7^ and STHdh^Q111/Q111^ cells were grown in Dulbecco’s modified Eagle’s medium (DMEM) supplemented with 10% fetal bovine serum (FBS) and 4 mM L-glutamine at 33 °C in a humidified atmosphere with 5% CO_2_.

### Gene knockout and overexpress

Gene knockouts of *p300*, *CBP*, *HDAC1*, *HDAC2, HDAC1/HDAC2* and *MSH3* were performed in HeLa or HeLa(CAG)_45_ using CRISPR-Cas9 technologies, as previously described ^70^. Briefly, HeLa or HeLa(CAG)_45_ cells were transiently transfected with desired Lenti-Crispr V2 plasmids using Lipofectamine 3000 reagent (Thermo Fisher Scientific Inc.) for 48 hours. Cells were then cultured in a 100-mm dish in medium containing 10 μg/μL puromycin. Cell clones were established from puromycin resistant cells, and screened for knockout by Western blotting analysis, and verified by DNA sequencing.

To establish HeLa-(CAG)_45_ *MSH3-*KO cells expressing various MSH3 isoforms (WT, FY-AA, ΔF2 and ΔF4), HeLa-(CAG)_45_ *MSH3-*KO cells were transduced with corresponding MSH3 Lentiviruses, which were produced by co-transfecting HEK293FT cells with the individual *MSH3* Lentiviral plasmids (plx-317-*MSH3*) and Lentiviral help vector (psPAX2, pMD2.G). The resulting HeLa-(CAG)_45_ *MSH3-*KO clones expressing individual MSH3 isoforms were screened with 10ug/ml puromycine (Sigma) and verified by Western blotting analysis for successful MSH3 rescue.

HeLa-(CAG)_45_, HEK293, STHdh^Q^^7/Q7^ and STHdh^Q111/Q111^ cells overexpressing Flag-HA-tagged H3WT, H3K56Q and H3K56R were transduced with corresponding Lentiviruses, which were produced by co-transfecting HEK293FT cells with the individual wild-type or mutant (K56Q and K56R) H3.1 or H3.3 Lentiviral plasmids (pCDH-CMV-*H3*) and Lentiviral help vector (psPAX2, pMD2.G). The resulting clones were screened with 10 µg/mL puromycine (Sigma) and verified by Western blotting analysis with anti-H3 and anti-Flag or anti-HA antibodies. Clones expressing equal levels of endogenous H3 and recombinant H3 were used for further analysis.

### Protein and nuclear extract preparation

MutSɑ and MutSβ were overexpressed in High Five insect cells using baculovrius system and purified as described ^21^. GST-MSH3, and GST-MSH6 fragments were expressed in the *Escherichia coli* strain BL21. Cultures were grown at 37 °C with shaking at 250 rpm. Protein expression was induced at OD_600_=0.4-0.6 by the addition of 0.5 mM IPTG at 18°C for 16-18 hours. GST-MSH6 domain II or GST-MSH3 fragments were purified as described ^71^. Briefly, cells were lysed in the lysis buffer (20 mM Tris HCl pH8.0, 150 mM NaCl, 0.1% NP-40, 1 mM EDTA, 1 mM DTT, and protease inhibitors) followed by sonication for 20 min and centrifugation (30 min at 12000 g). Supernatants were chromatographed through FPLC columns GSTrap and Superdex 200. The purified proteins were stored in aliquot at -80 °C. The nuclear extracts were prepared as described ^72^.

### Protein pulldown assays

Biotinylated peptides (H3K9, H3K9ac, H3K36, H3K36me3, H3K56, H3K56ac and H3K56me1) were synthesized at ABclonal Technology (Woburn, MA). Peptide pulldown assay was performed as described previously ^73^. Briefly, cell extracts or purified proteins were incubated with 10 µg of peptides prebound to streptavidin agarose beads (Pierce) overnight at 4°C in the binding buffer (20 mM HEPES pH 7.9, 150 mM KCl, 0.1% Triton X-100, 20% v/v glycerol, 0.2 mM EDTA and protease inhibitors). The mixture was washed six times with the washing buffer (20 mM HEPES pH 7.9, 300 mM KCl, 0.1% Triton X-100, 20% v/v glycerol, 0.2 mM EDTA and protease inhibitors). The bound proteins were analyzed by Western blotting analysis after gel electrophoresis.

Nucleosome pulldown assay was performed as described previously ^74^. Briefly, nucleosomes (2 µg) were incubated with 600 ng of purified MutSɑ or MutSβ at 4°C in the binding buffer (50 mM Tris pH 7.5, 0.5% NP40, 200 mM NaCl, protease inhibitors) for 3 hours. Streptavidin resin slurry (40 µL, Invitrogen) was added to the mixture for further incubation overnight at 4°C. Streptavidin beads were recovered and washed 6 times with the washing buffer (50 mM Tris pH 7.5, 0.5% NP40, 300 mM NaCl and protease inhibitors). The bound proteins were analyzed by SDS-PAGE and Western blotting analysis.

### Co-immunoprecipitation and Western blotting analysis

Nuclear extracts (1 mg) were incubated with 3 μg of α-IgG (Santa Cruz Biotechnology), α-H3 antibody (Abcam) in binding buffer (20 mM Tris pH 8.0, 137 mM NaCl, 1% NP-40, 2 mM EDTA, 1 mM DTT and protease inhibitors) at 4 °C for 3 hours. 40 μL agarose-protein G beads (50% slurry; Invitrogen) were pre-equilibrated with a buffer containing 20 mM Tris pH 8.0, 137 mM NaCl, 1% NP-40, 2 mM EDTA, 50 μg/ml bovine serum albumin, 1 mM DTT and protease inhibitors. The equilibrated beads were added into the reaction mixture, and incubated with rotation at 4 °C overnight. The beads were washed three times with a high salt washing buffer (10 mM Tris pH 7.4, 1 mM EDTA, 1 mM EGTA, 500 mM NaCl, 1.0% Triton X-100 and protease inhibitors) and three times with a low salt washing buffer (10 mM Tris pH 7.4, 1 mM EDTA, 1 mM EGTA, 150 mM NaCl, 1.0% Triton X-100 and protease inhibitors). The bound proteins were analyzed by SDS-PAGE and Western blotting analysis.

### Chromatin fraction assay

Preparation of chromatin fraction was performed as described previously ^75^. Briefly, cells pellets were washed with cold PBS buffer and subsequently re-suspended in the lysis buffer containing 10 mM HEPES pH 7.4, 10 mM KCl, 0.05% NP-40 and protease inhibitors, and incubated on ice for 20 minutes. Cell mixtures were separated by centrifugation (15,000 rpm for 10 min) at 4°C, and the supernatant was kept for cytoplasm fraction. After wishing twice with the lysis buffer, nuclei pellets were re-suspended in the low salt buffer (10 mM Tris HCl pH 7.4, 0.2 mM MgCl_2_, 1 % Triton X-100 and protease inhibitors) and incubated on ice for 15 min, followed by centrifugation (15,000 rpm for 10 min) at 4°C. The supernatant contained the nucleoplasm proteins, and the pellet contained the chromatin, which was re-suspended in 0.2 N HCl and incubated on ice for 20 min. After centrifugation (15,000 rpm for 10 min), the supernatant was neutralized with 1 M Tris-HCl (pH 8.0) and defined as the chromatin fraction.

### In situ proximity ligation assay

In situ proximity ligation assay (PLA) in combination with immunofluorescence confocal microscopy was performed to detect the H3-MutSβ interaction. Briefly, the fixed cells (HEK293 3.1 WT, H3K56Q and H3K56R) were incubated with primary antibodies: mouse monoclonal antibody against MSH3 and rabbit polyclonal antibody against Flag. Oligonucleotide-conjugated probe secondary antibodies (PLA PLUS and MINUS) against the primary antibodies initiate a DNA amplification reaction to generate signals that depend on the close proximity of two proteins (<40 nm).

### Immunofluorescence analysis

Preparation of chromatin-bound protein in immunofluorescence (IF) were described previously with some modifications ^49,50^. Briefly, cells were washed with the CSK buffer (10 mM PIPES/KOH pH 6.8, 100 mM NaCl, 300 mM Sucrose, 1 mM EGTA, 1 mM MgCl2, 1 mM DTT, 0.5% Triton X-100 and protease inhibitors) once and then pre-extracted with cold freshly made CSK/0.5% Triton X-100 buffer at 4°C for 20 min. Cells were fixed with 4% paraformaldehyde and permeabilized with 0.2% Triton X-100 in PBS. The fixed cells were blocked by 5% (w/v) bovine serum albumin (BSA/PBS) at room temperature for 1 hour, followed by incubating with a primary antibody at 4 °C overnight. After washing with PBS 3 times (5 minutes each), the cells were incubated with secondary antibody (anti-rabbit Alexa Fluor 488 and anti-mouse Alexa Fluor 555) for 2 hours at room temperature. Cells were mounted in ProLong GOLD antifade reagent with DAPI (Life Technologies), and images were determined by a Leica DMi8 confocal scanning laser microscopy system.

### PCR, genotyping and *HTT* CAG repeat analysis

The CAG repeat analysis for HeLa (CAG)_45_ cells was performed as described ^54^ previously. Briefly, genomic DNA (100 ng) was used for PCR reaction under the following conditions: 94°C (5 min); 35 cycles of 94°C (30s), 55°C (30s), 72°C (2 min); and 72°C (7 min). PCR products were resolved in a 1.5% agarose gel, and visualized by GelGreen staining.

The *HTT* CAG repeat analysis in STHdh^Q7/Q7^ and STHdh^Q111/Q111^ cells were performed as following. The *HTT* CAG repeats were analyzed by PCR using primers specific for the *HTT* CAG repeats from knock-in allele but not for the mouse sequence. PCR reaction conditions were 94°C (4 min); 40 cycles of 94°C (45s), 68°C (45s), 72°C (4 min); and 68°C (1 min) 72°C (10 min). The forward primer was fluorescently labeled with 6-FAM and products were resolved using ABI 3730xl DNA analyzer (Applied Biosystems) with GeneScan 500 ROX as internal size standard (Applied Biosystems). GeneMapper v3.7 (Applied Biosystems) was used to generate CAG repeat size distribution traces.

## Acknowledgments

The work was supported in part by a grant from Chinese Institutes for Medical Research (to GML) and a National Institutes of Health intramural research grant (DK036119, to WY). We thank Dr. Liya Gu for stimulating discussion and technical help.

## Author Contributions

Conceptualization: GML and WY

Methodology: JG, WY, and GML

Investigation: JG, WY, and GML

Writing – original draft: JG and GML; review & editing: JG, GML and WY

## Data and materials availability

All data are available in the main text or the supplementary materials.

