## Supplemental Figures for "Interaction between histone residue H3K56 and mismatch repair protein MutSβ drives CAG repeat expansion"

**Supplemental Data**

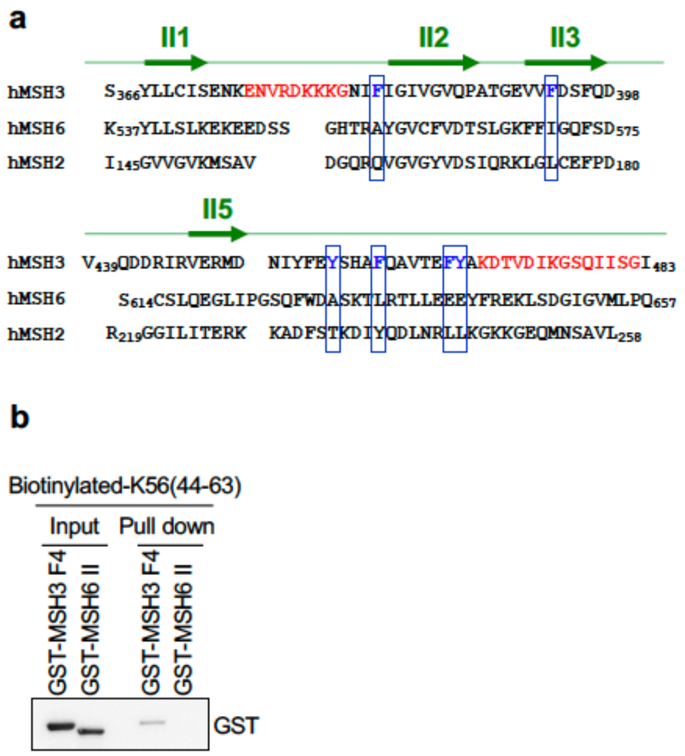


**Figure S1. MSH3 domain II and its interactions with H3 peptides. a,** Comparison of amino acid sequences of domain IIs between human MSH3, MSH6 and MSH2. Red-typed amino acid residues indicate the disordered regions, and blue typed residues indicate the critical aromatic residues in MSH3, which are structurally close to the disordered regions. Blue columns show aromatic residues in human MSH3 and their corresponding residues in MSH6 and MSH2. **b,** Peptide pulldown assay showing interactions between biotinylated-K56(44-63) peptide and GST-MSH3-F4 (MSH3 Domain II) and GST-MSH6 Domain II.

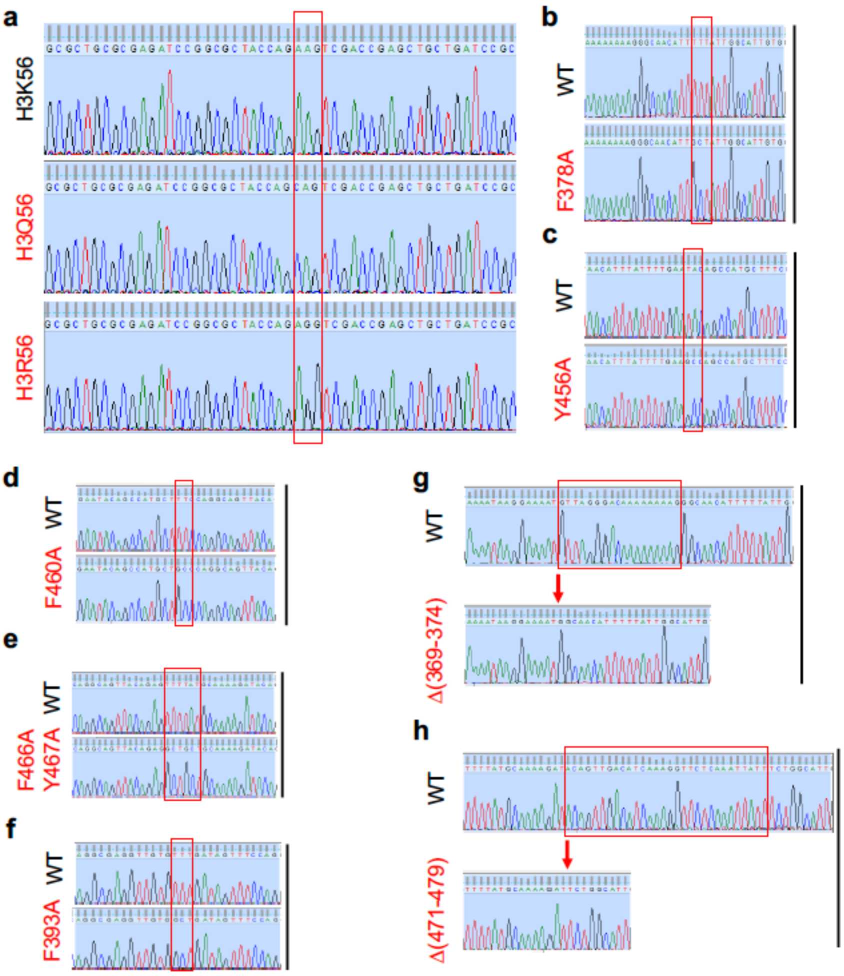


**Figure S2. DNA sequencing analysis verifying site-specific mutagenesis.** Sequencing results for H3K56, H3K56Q and H3K56R (**a**), MSH3 aromatic WT and MSH3-F378A (**b**), MSH3-WT and MSH3-Y456A (**c**), MSH3-WT and MSH3-F460A (**d**), MSH3-WT and MSH3-F466A-Y467A (**e**), MSH3-WT and MSH3-F393A (**f**), MSH3-WT and MSH3-Δ(369-374) (**g**), MSH3-WT and MSH3-Δ(471-479) (**h**)

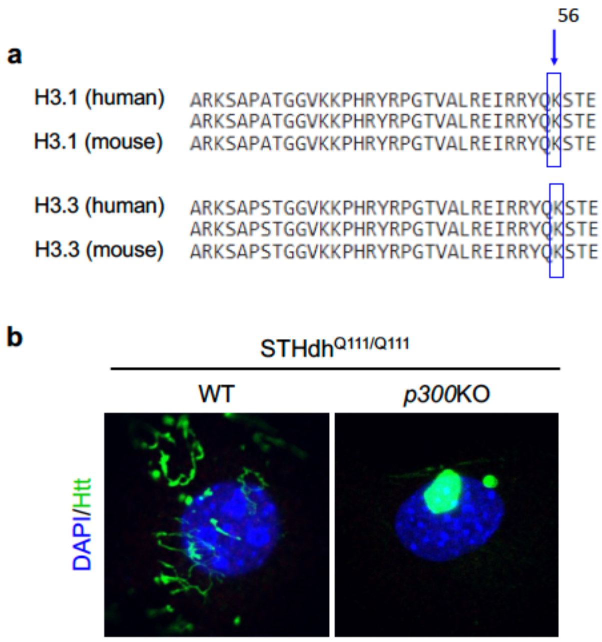


**Figure S3. a,** Comparison of amino acid sequences between H3.1 (human) and H3.1 (mouse), as well as H3.3 (human) and H3.3 (mouse). The blue column and arrow indicate the conservation of K56 between human and mouse H3. **b**, Representative immunofluorescence images showing Htt staining in WT and *p300*-KO STHdh$^{Q111/Q111}$ cells. WT and *p300*-KO STHdh$^{Q111/Q111}$ cells were culturing for 18 weeks.
